# The remarkable similarity in the serum proteome between type 2 diabetics and controls

**DOI:** 10.1101/2024.03.19.585746

**Authors:** David E. Mosedale, Tilly Sharp, Adam de Graff, David J. Grainger

**Affiliations:** Methuselah Health UK Ltd. B950 Dorothy Hodgkin Building Babraham Research Campus Cambridge, CB22 3FH

**Author notes:** Corresponding author: DEM. Currently at: RxCelerate Ltd., B950 Dorothy Hodgkin Building, Babraham Research Campus, Cambridge CB22 3FH (David E. Mosedale, Tilly Sharp, David J. Grainger). Currently at: InVitro Cell Research LLC, Greater New York Metropolitan Area, USA Word count: 4400. Four figures and two tables. Abbreviations – T2DM: type 2 diabetes mellitus, PTM: post-translational modification, IGF: Insulin-like growth factor, MS: mass spectrometry, ALS: IGF-binding protein complex acid-labile subunit, IGFBP: IGF binding protein.

## Abstract

Type 2 diabetes mellitus (T2DM) is a rapidly increasing threat to global health, which brings with it a demand for better treatments. This study aimed to identify differences in the proteome of patients with T2DM to identify new targets for therapeutic intervention. We used a highly reproducible bottom-up proteomics protocol to investigate differences in protein, peptide and post-translational modifications between subjects with T2DM and matched controls in an untargeted manner. The serum proteome was remarkably similar at the protein level with no differences between the subject groups across 175 proteins and five orders of magnitude. Strong associations were found, however, between fasting serum glucose levels and glycations of abundant serum proteins, including sites on apolipoprotein A1, apolipoprotein A2 and α2- macroglobulin. We also investigated proteome differences associated with BMI, and found all three components of the ternary complex (IGF-binding protein complex acid-labile subunit (ALS), IGF-binding protein 3 (IGFBP-3) and IGF-2) were strongly negatively associated with BMI. The results show the power of a proteomics protocol optimised for precision rather than depth of coverage, which here has identified strong correlations between physiological measurements and very low abundance post-translational modifications. In T2DM any differences in the serum proteome are very small, and likely a consequence rather than a cause of hyperglycaemia.

**Article highlights:**

- Our goal was to use high-precision label-free bottom-up LC-MS/MS proteomics to investigate differences in the proteome of patients with T2DM and controls, and potentially identify novel targets for future research.
- The serum proteome is remarkably similar in patients with T2DM and controls, with the only major difference being glycations of abundant serum proteins
- All three components of the ternary complex (comprised of ALS, IGFBP-3 and IGF-2) were strongly negatively associated with BMI.
- The results highlight the power of a proteomics study designed with three key features at its core: a proteomics protocol optimised for precision rather than depth of coverage; an open bioinformatics approach investigating proteins, peptides and PTMs without prior assumptions about which features are important; and analysis of individual subject samples so that results take into account person-to-person variability

## Introduction

Type 2 diabetes mellitus (T2DM) is a multi-factorial metabolic disease, characterised by high blood glucose and insulin resistance. The pancreas in patients with T2DM continues at least at first to produce insulin, yet its target tissues become resistant to its action. Some individuals maintain a minor insulin resistance, while others progress to manifest T2DM (1), which has a variety of acute and chronic consequences, including increased thirst and volume of urination, hyperglycemia, and microvascular and macrovascular complications.

Fortunately, the onset of T2DM can be delayed or prevented through simple lifestyle changes. However, despite the well-known benefits of exercise and prevention of obesity, the clinical incidence of T2DM continues to grow (2,3). The most effective first-line treatment for T2DM is metformin. However, in a situation analogous to the disease itself, a wide range of effects of metformin have been catalogued, while the molecular mechanism of action of metformin remains incompletely understood (4).

The pathogenesis of T2DM at the molecular level involves complex interactions between genetics and the environment. However, while there is a substantial contribution of genetics to T2DM (heritability estimates vary from 25 to 72%) relatively little of this has been explained by identification of over 100 genetic loci associated with the disease (5). There is a substantial knowledge gap here, which large genome-wide association studies carried out by international consortia have failed to resolve. This has led to T2DM being the target of many proteomics analyses over the years, trying to uncover patterns of protein expression that lead to insights into the pathogenesis of the disease. Early studies used 2D gel electrophoresis and mass spectrometry (MS) to identify proteins with differential expression in serum or plasma between T2DM and controls (6–16). These studies identified a wide range of biomarkers, few of which were found consistently across the different studies. The intrinsic low throughput of this technique limits the power of these studies, with most analysing just a few pooled samples.

More recently, multiplexed protein assays have been used to identify patterns of protein expression in patients with T2DM using a variety of techniques (17–21). SELDI or MALDI-TOF MS have also been applied to the analysis of T2DM with some success (22–25). None of the above techniques, however, lend themselves to the analysis of post-translational modifications (PTMs) of proteins that may be associated with the pathogenesis of the disease.

Bottom-up proteomics studies, which offer the best non-hypothesis driven approach to investigating both proteins and PTMs, have been carried out on patients with T2DM (26–29). However, of these studies, samples were often pooled prior to analysis, used limited number of samples or in one case was carried out on an unusual matrix (tears). Pooling of samples has a distinct disadvantage in that it removes person-to-person variation from the data, leading to bias in the results.

We have extensively optimised bottom-up proteomics protocols to improve reproducibility of the results and have applied this method to individual serum samples from patients with T2DM and matched controls. The protocol to identify peptides was designed to include the possibility of identifying a wide range of post-translational modifications as potential biomarkers between the two groups of patients.

## Research design and methods

### Study subjects

The samples used for this study were selected from among the 1334 subjects recruited into the Metabolomics and Genomics in Coronary Artery Disease (MaGiCAD) study designed to study risk factors for coronary heart disease and carried out at Papworth Hospital NHS Trust, Cambridgeshire UK. Patients attended for a diagnostic angiogram, and were recruited into the study prior to undergoing the procedure. The MaGiCAD study was completed and all samples stored at -80°C until use in this proteomics study.

Samples were selected from among the male patients with a clinical diagnosis of T2DM. 16 patients were selected and matched on the basis of age, BMI and cardiovascular disease status with other patients from the MaGiCAD study with no diagnosis of diabetes. Wherever possible, patients with as few medications and concomitant diseases were chosen. No patients in the study were taking insulin. A summary of the study subjects is given in Table 1.

**Table 1.** Study subjects summary. In each cell of the table data is shown as mean +/- SD (95% confidence intervals), with the result of the paired t-test analysis for controls vs diabetics shown in the right hand column.

|  | Diabetic subjects | Controls | Paired t-test |
| --- | --- | --- | --- |
| Count | 16 | 16 |  |
| Age (years) | 61.4 +/- 5.1<br>(58.6 – 64.1) | 61.0 +/- 5.0<br>(58.3 – 63.7) | 0.232 |
| Sex (M/F) | 16 / 0 | 16 / 0 | n/a |
| BMI (kg/m<sup>2</sup>) | 28.8 +/- 4.8<br>(26.2 – 31.4) | 28.9 +/- 3.7<br>(27.0 – 30.9) | 0.842 |
| Major diseased coronary arteries | 1.5 +/- 1.2<br>(0.9 – 2.1) | 1.5 +/- 1.1<br>(0.8 – 2.1) | 0.334 |
| Fasting serum glucose (mM) | 10.9 +/- 5.1<br>(8.2 – 13.6) | 5.3 +/- 0.9<br>(4.8 – 5.8) | 0.0008 |

### Sample collection

Patients attended hospital on the morning of their angiogram following an overnight fast. Blood samples were taken from the arterial sheath during the angiogram but prior to injection of the contrast medium. Arterial blood was transferred to a 15 ml polypropylene tube and left to clot at room temperature for between 2 and 3 hours. The sample was then spun at 4000 g for 5 minutes and the serum removed. The serum was re-spun to pellet any remaining red blood cells and the supernatant taken and frozen in aliquots at -80°C until use.

### Preparation of sample for LC-MS/MS

Seven of the most abundant serum proteins were removed from the serum using a MARS7 spin cartridge (Agilent #5188-6408), following the manufacturer’s instructions. The column binds and removes >95% of albumin, IgG, IgA, transferrin, haptoglobin, α1-antitrypsin, and fibrinogen. The eluent was concentrated using Vivaspin 500 centrifugal concentrators (5 kDa molecular weight cut-off) that were washed before use and the protein concentration measured using a bicinchoninic acid protein assay kit (Pierce #23227).

Prior to LC-MS/MS analysis, samples were reduced, alkylated and digested with trypsin using the In-Solution Tryptic Digestion and Guanidination Kit from Thermo Scientific (#89885). Briefly, 10 µg of protein was incubated with 5 mM DTT (Thermo #20291) in 50 mM ammonium bicarbonate buffer (Sigma #A6141) for 5 minutes at 95°C. The sample was cooled for 20 minutes and iodoacetamide (Thermo #90034) added to 10 mM before incubation in the dark for 30 minutes. Samples were buffer-exchanged to 50 mM triethylammonium bicarbonate buffer (Thermo #90114) by diluting and concentrating using Vivaspin concentrators (three repeats of approximately 20x dilution), concentrated to approximately 30 µl and 1 µl of trypsin (Pierce trypsin protease, MS grade; Thermo #90057) added and incubated for 3 hours at 37°C. An additional 1 µl of trypsin was added to the reaction and the samples incubated for 16 hours at 30°C. Samples were then frozen until LC-MS/MS analysis.

### LC-MS/MS

LC-MS/MS was performed using a Dionex Ultimate 3000 RSLC nanoUPLC system and a QExactive Orbitrap mass spectrometer. Separation of peptides was performed by reverse-phase chromatography at a flow rate of 300 nL/min and a Thermo Scientific reverse-phase nano Easy-spray column (Thermo Scientific PepMap C18, 2μm particle size, 100A pore size, 75μm i.d. x 50cm length). Peptides were loaded onto a pre-column (Thermo Scientific PepMap 100 C18, 5μm particle size, 100A pore size, 300μm i.d. x 5mm length) from the Ultimate 3000 autosampler with 0.1% formic acid for 3 minutes at a flow rate of 10 μL/min. After this period, the column valve was switched to allow elution of peptides from the pre-column onto the analytical column. Solvent A was water + 0.1% formic acid and solvent B was 80% acetonitrile, 20% water + 0.1% formic acid. The linear gradient employed was 2-40% B in 90 minutes (120 minute total run time).

The LC eluent was sprayed into the mass spectrometer by means of an Easy-spray source (Thermo). All m/z values of eluting ions were measured in an Orbitrap mass analyzer, set at a resolution of 70000. Data-dependent scans (Top 20) were employed to automatically isolate and generate fragment ions by higher energy collisional dissociation (HCD) in the quadrupole mass analyser with dynamic exclusion not employed. Measurement of the resulting fragment ions was performed in the Orbitrap analyser, set at a resolution of 17500. Peptide ions with charge states of 2+ and above were selected for fragmentation.

### Data analysis and bioinformatics

Progenesis QI for Proteomics v4.2 (Nonlinear Dynamics, Waters) was used for peak quantification. Briefly, all 32 samples were aligned and peak picking carried out. One sample was found to give poor results and it and its matched pair were excluded from further analysis. Ions with a charge of 1 were excluded, and MS data deisotoped before being exported for ion identification.

Ion identification in Mascot v2.8 (Matrix Science) was carried out by combining two searches (a narrower, more robust search and a wider error tolerant search), to gather putative identifications on as many peptides as possible. Ten MS2 scans per peak were searched against two protein databases– SwissProt restricted to human proteins and the common Repository of Adventitious Proteins (cRAP). Common settings were use of the enzyme trypsin, allowing up to 3 missed cleavages, no fixed modifications, peptide mass tolerance +/- 5 ppm, MS/MS tolerance +/- 0.02 Da. Both searches were controlled by searching a decoy database with the PSM false discovery rate set at 1%. The first search was carried out with nine variable modifications (carbamidomethyl (C), deamidation (NQ), oxidation (M), cation:Na (DE), cation:Fe[III] (DE), cation:Ca[II] (DE), gln -> pyro-glu (N-term Q), pyro-carbamidomethyl (N-termC) and Cation:Mg[II] (DE)) This list of nine variable modifications were the most common modification types found in an initial survey error tolerant search carried out with just a single variable modification (carbamidomethyl (C)) but with the full Unimod database of modification types available to Mascot.

The second search was an error tolerant search, with three variable modifications (carbamidomethyl (C), deamidation (NQ) and oxidation (M)) but with the full Unimod database of modification types available to Mascot. In order to reduce the number of unsound identifications the scores obtained in this second search were all reduced by a factor of 100, before combining both sets of results within the Progenesis software. This score reduction enabled results from the two sets of searches to be distinguished.

A list of all of the identifications obtained from both searches was exported from Progenesis and where a single deconvolved ion peak was given more than one identity resolution of the conflict was carried out using a simple algorithm built from the results of manually resolving conflicts on six full datasets by two operators.

Following conflict resolution any peptide that could have arisen from more than one protein was excluded from the peptide list. Protein abundances were calculated by the sum of all remaining peptides (all now unique to their parent protein). Post-translational modification (PTM) fractions were calculated using a custom script (Bio::Neos) that for every identified unique modification / site calculates the sum of the abundance of all peptides containing the modification and divides it by the sum of the abundance of all peptides containing that amino acid (whether modified or not) generating a number between 0 and 1 representing the fractional modification for every modification / location / sample combination. The MS2 fragmentation patterns of key peptides containing modifications of interest were reviewed manually to confirm identity.

Statistical analyses were carried out in R (v4.2.0) and R studio (v 2022.02.3). False discovery rate (FDR) correction was carried out in GraphPad Prism (v9.4.0) using the two-stage step-up method of Benjamini, Krieger and Yekutieli (30) with a desired FDR of 20%.

### Data and resource availability

The data generated during the current study are available from the corresponding author upon reasonable request.

## Results

### The serum proteome is remarkably similar in patients with T2DM and controls

The LC-MS/MS protocol we used for this proteomics study was carefully optimised to reduce sample-to-sample variability in the measurement of the abundances of the individual peptides, not to increase the depth of coverage or number of proteins measured. Using this protocol we identified 4855 peptides from 213 proteins. However, all of the analyses described below were restricted to the 4817 peptides from 175 proteins with at least two peptides uniquely derived from the parent protein.

We began by carrying out a paired analysis of protein abundances between the control and diabetic subjects. While previous proteomics studies have often found numerous differences between patients with diabetes and controls, we found that the total protein abundances of the 175 proteins with at least two unique peptides were remarkably similar between the controls and the patients (Figures 1A and 1B). Across all of the proteins none had an unadjusted p value < 0.01 for being different between the controls and the diabetics (paired t-test), and none were significant following false discovery rate correction at 20%.

**Figure 1.**
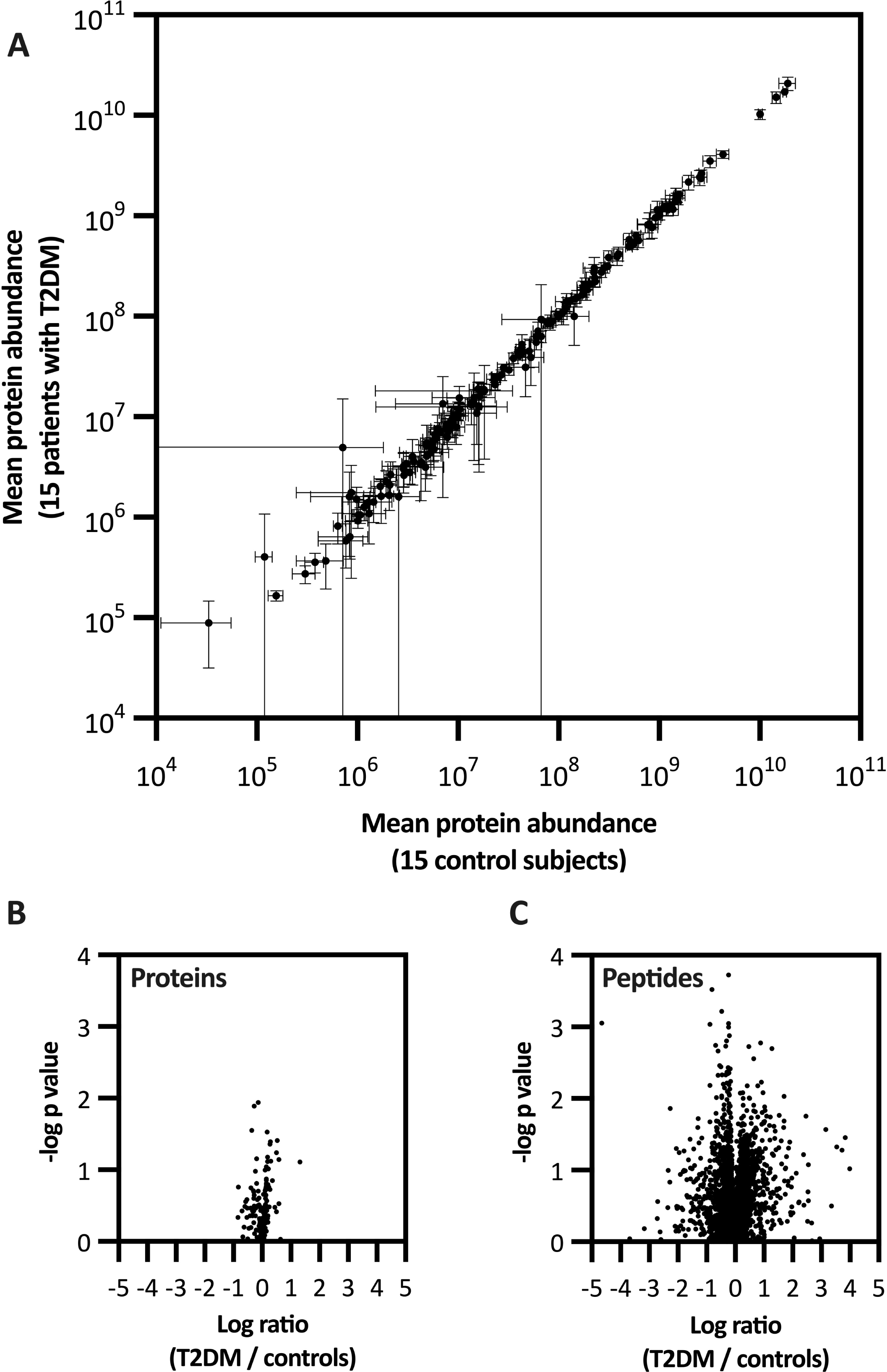
Extreme similarity in protein abundances in serum from patients with T2DM and matched controls. Figure 1A shows data for each protein with the average abundance of each protein for the 15 patients with T2DM plotted against the average abundance for the 15 control subjects. Mean data is plotted +/- 95% confidence intervals. Figures 1B and 1C show volcano plots of the -log p values for paired analysis of the proteins (Fig 1B) and peptides (Fig 1C) vs. the mean ratio of the proteins and peptides abundances respectively (T2DM / controls).

While our initial analysis showed no difference between patients with T2DM and controls we also examined the individual peptides for differences between the two groups, to see if there were differences that could be isolated to individual protein domains or fragments. In total 4817 peptides from the proteins with at least two unique peptides were analysed and while inevitably with such a large dataset some peptides were found to be different at p < 0.01 (50/4817; 1.04%; paired t-test), none were outliers in the volcano plot (Figure 1C) nor significant following correction using false discovery rate at 20%.

### Associations between glucose and the serum proteome

Separately to the paired analysis between the diabetic patients and control subjects we analysed the correlations between fasting plasma glucose concentrations and the serum proteome. One of these associations was markedly stronger than the others: serotransferrin showed a negative correlation (r = -0.647, p = 1.12E-04) with glucose. However, serotransferrin is one of the seven proteins removed by the MARS7 spin cartridge which we used to remove the most abundant serum proteins prior to LC-MS analysis. It remains to be determined whether there is a negative association between glucose serotransferrin abundance in serum prior to application on to the column or whether there is differential removal of serotransferrin in serum samples containing varying levels of glucose. In addition to serotransferrin the association between serum glucose and a further four proteins survived correction using FDR: protein z-dependent protease inhibitor, complement C1q subcomponent subunit C, biotinidase and cartilage oligomeric matrix protein (Figure 2) but all four were of borderline significance (q values between 0.1 and 0.2).

**Figure 2.**
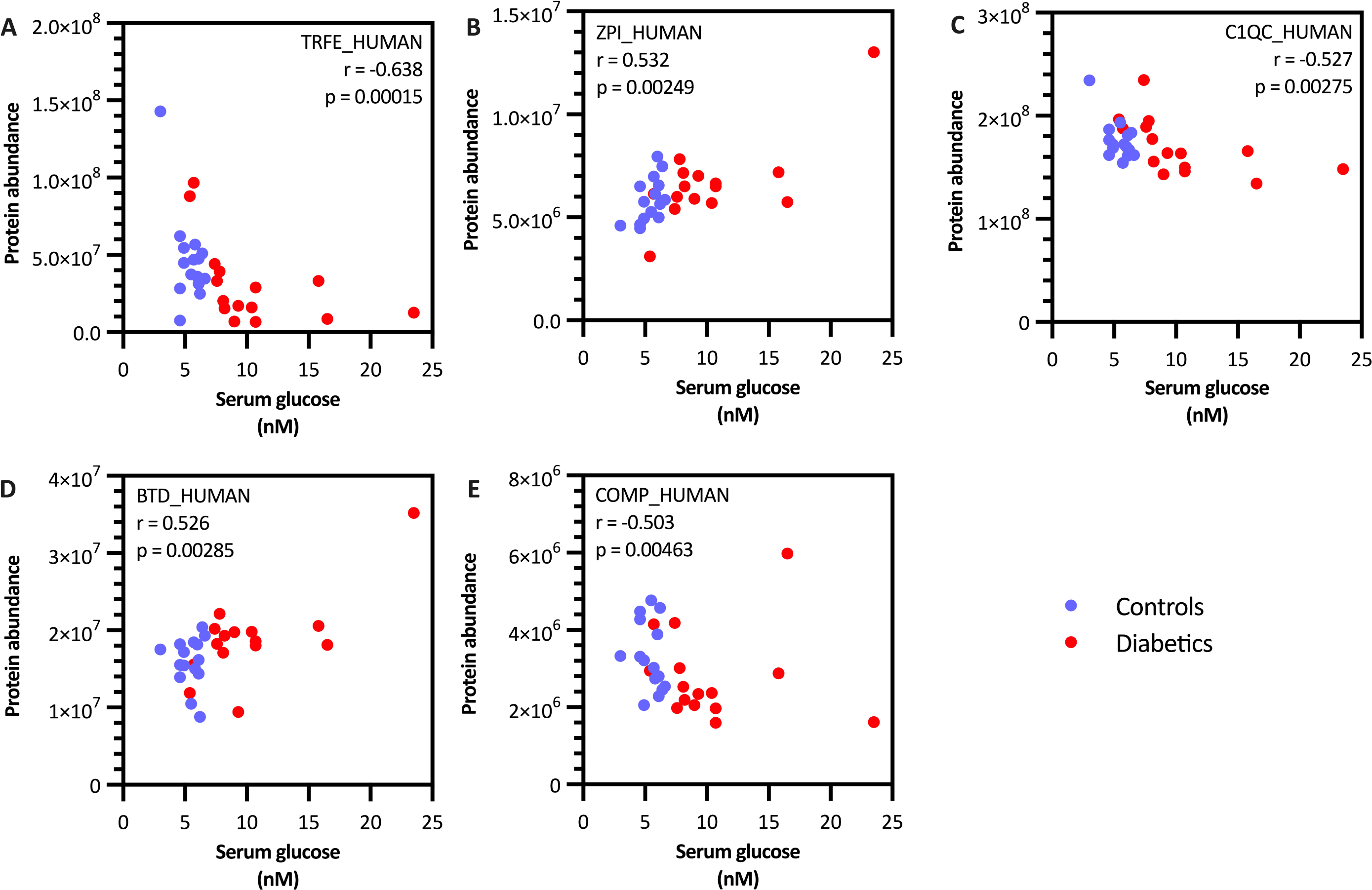
Correlations between serum fasting glucose and five serum proteins that remain significant after adjusting using FDR at 20%. Data from control subjects shown in blue and patients with diabetes in red. Spearman’s rank correlation coefficient and unadjusted p values are shown in the upper right of each graph. A Serotransferrin; B Protein Z-dependent protease inhibitor; C Complement C1q subcomponent subunit C; D Biotinidase; E Cartilage oligomeric matrix protein.

We proceeded to look at the association of the 4817 individual peptides with glucose. 33 peptides correlated with glucose at p < 0.001, all of which plus another 6 peptides remained significant following FDR correction at 20% (Supplementary Table 1, sorted by protein). These fall into three main categories. The largest category, with 20 significant negative correlations, were correlations with peptides from serotransferrin, which at the protein level we saw strongly negatively correlated with glucose concentrations. Secondly, we found four peptides from apolipoproteins A1 & A2 with a hex-modification that show a strong positive correlation with glucose. These are likely to be direct consequences of the high glucose concentrations. Thirdly, we found significant negative correlations with eight peptides from complement C3. There was a tendency for peptides from complement C3 to correlate negatively with glucose (median Spearman’s correlation coefficient was -0.13), but protein abundance of complement C3 did not correlate with glucose nor was there an association between location across the protein and correlation with glucose (data not shown). Fourthly, we have seven peptides, one each from seven further proteins, that pass FDR correction for correlation with glucose.

### Post-translational modifications

The peptide abundances were used to calculate PTM fractions which are a number between 0 and 1 representing the fractional modification for every modification / location / sample combination. We tentatively identified 1463 PTMs in our data, of 179 distinct types, after restricting to modifications whose underlying residue was also found on at least one other peptide without the modification of interest. The search strategy used to identify peptides was deliberately open to finding any of the modifications in Unimod. As a consequence, due to the large number of MS2 scans searched, we would expect the results to contain some peptide identities that would not be correct, even when results are limited to 1% FDR. Indeed, we found that just over half (103; 58%) of the distinct types of PTM in our data were found just once and were probably spurious. Nevertheless 83% of all the modifications identified (1215) were of a type found on at least 5 occasions, and all modifications were included in the following analysis.

The PTM fractions of two modifications were significantly different between T2DM and their paired controls with an unadjusted p value < 0.001 (paired t-test), the most significant of which survived FDR correction at 20%. Both of these PTMs were hexose modifications of lysine residues in α2-macroglobulin. Moreover, the next three most significant differences were increases in hexose modifications in α2-macroglobulin and apolipoprotein A1 in patients with T2DM (not shown). We also examined the correlation of the PTM fractions with glucose, and found 6 correlations strongly significant (p < 0.001), of which the four most significant remained significant after FDR correction at 20% (Figure 3 and Table 2). All of these were also hexose modifications of lysine residues, and included the two modifications that were different between T2DM and their paired controls.

**Figure 3.**
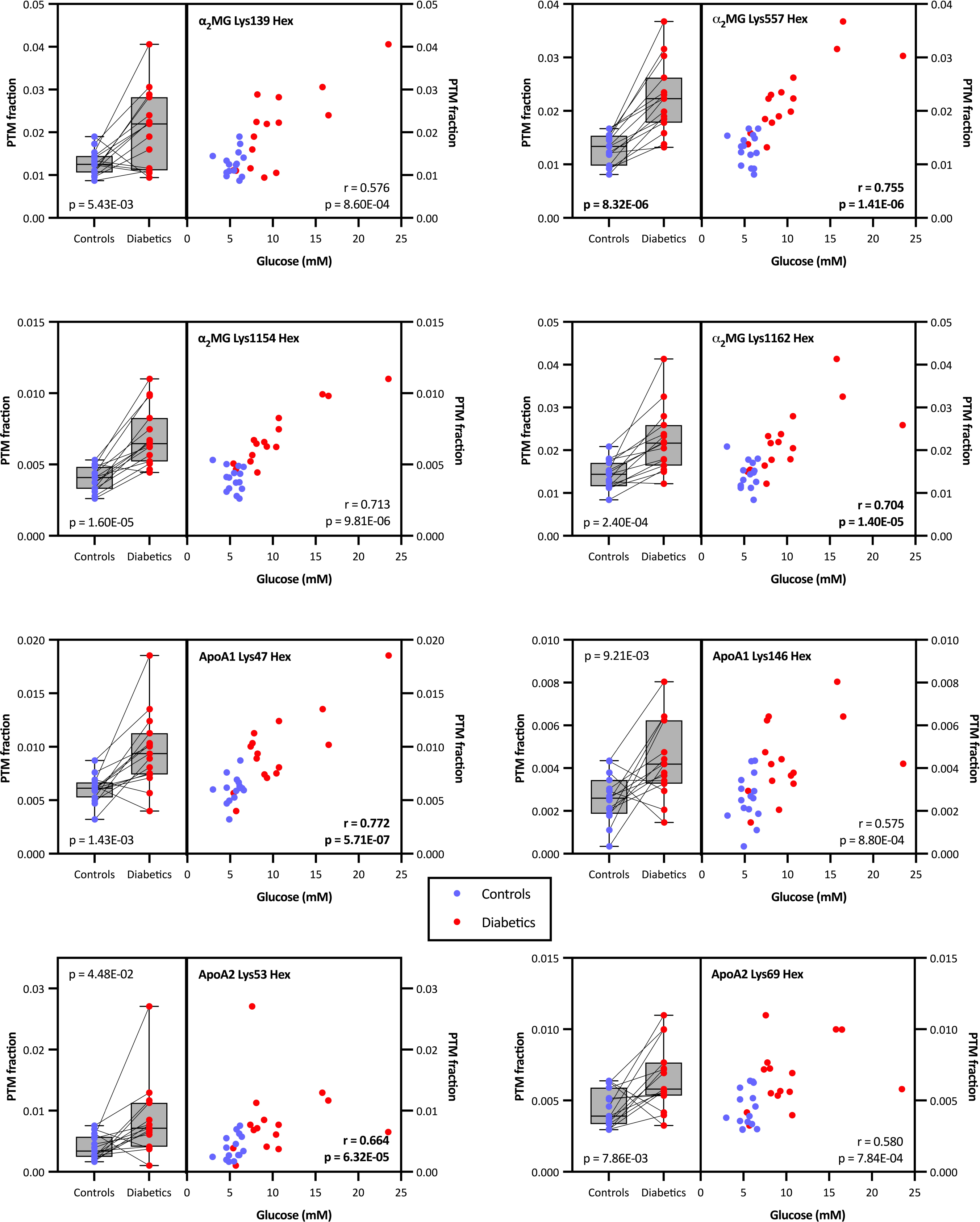
Associations between hexose modifications and patient group and fasting glucose. Data shown are all of the PTMs that were p < 0.001 (unadjusted) in either the non-hypothesis or hypothesis-driven analyses for the paired t-test for patients with T2DM vs controls or the Spearman’s correlation coefficient with fasting glucose. Each of the eight graphs shows the same data represented in two ways: on the left controls vs diabetics, with lines joining the patients with diabetes and their respective control; on the right, where the same PTM fractions are plotted vs the fasting glucose concentrations. Data from control subjects shown in blue and patients with diabetes in red. Unadjusted p values are shown in each plot, with those that remained significant after adjustment for multiple testing using FDR shown in bold.

**Table 2.**
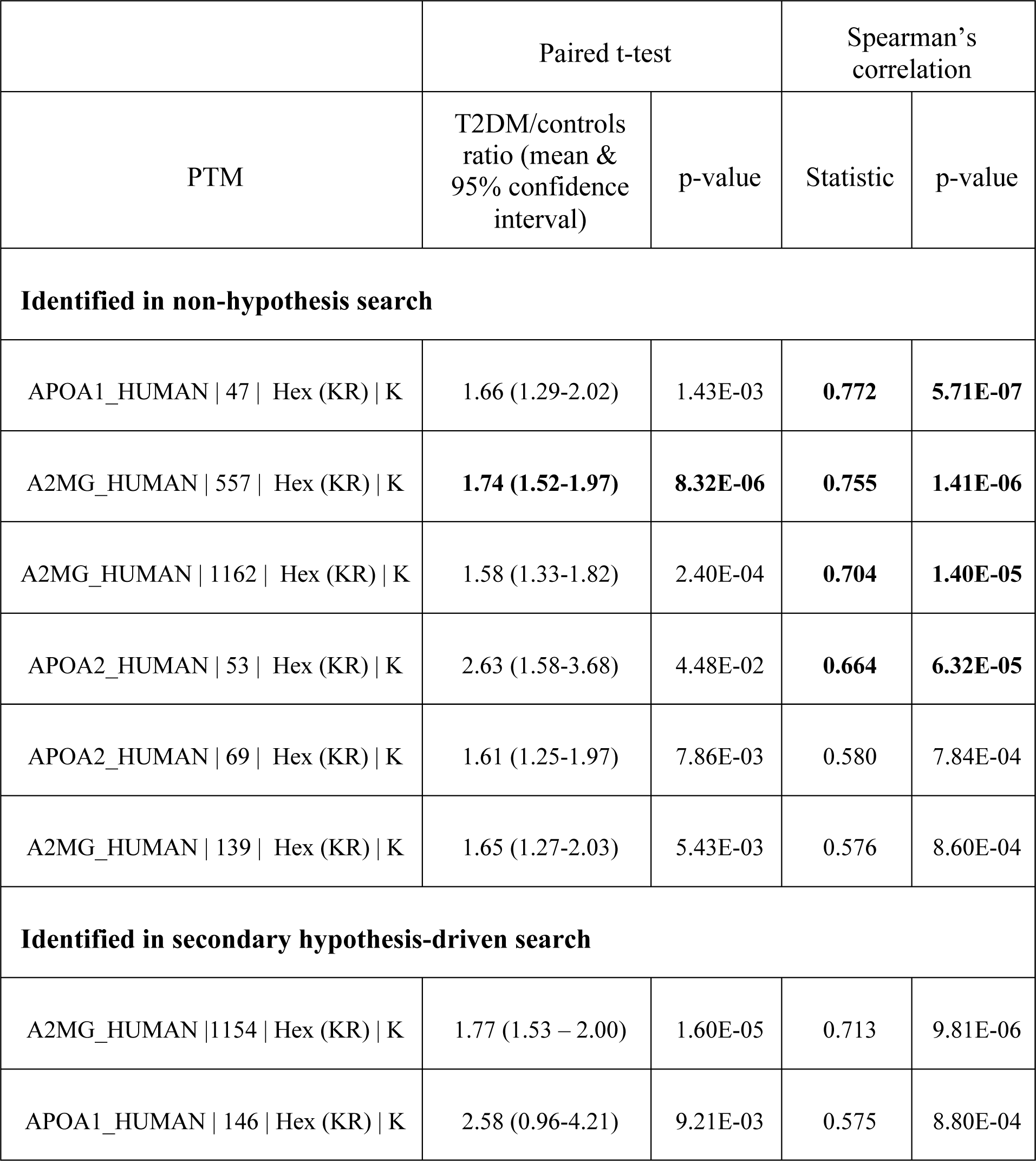
PTM fractions found associated with T2DM and correlating with glucose concentrations. All PTM fractions that were identified in either the primary non-hypothesis analysis or the secondary hypothesis-driven analysis and found to associate with T2DM at p < 0.001 using a paired t-test or correlating at p < 0.001 with glucose are shown in the table. Those associations identified in the non-hypothesis search that remain significant after FDR correction at 20% are shown in bold.

### Glycations

In total, in the untargeted analysis carried out we found 13 hexose modifications or glycations of common serum proteins, six of which correlate strongly with serum glucose. The MS2 fragmentation patterns of all of the hex-modified peptides were checked for accuracy and confirmed. We reran the MS identification searches to see if we could identify any further glycation sites, by including Hexose (Unimod #41, + 162.052824) in both the standard and error tolerant searches as an additional variable modification. The same 13 modifications found in the original untargeted search were identified again, plus an additional 31 glycations. In general, however, these were not good identifications, with manual review of the MS2 fragmentation patterns confirming only five of these additional modifications. All five of the additional validated hex modifications correlated positively with glucose at p < 0.05, however, with two at p < 0.001 (Table 2).

### The serum proteome and obesity

In addition to the core comparisons between the serum proteome and T2DM we extended our analysis to compare the serum proteome with BMI, a key risk factor in T2DM. While the patients with T2DM and controls were matched on the basis of BMI, subject BMIs ranged from 19.3 to 36.9 across all subjects.

Seven of the proteins showed Spearman’s correlations with statistical significance of < 0.01, of which three remained significant after false discovery rate correction at 20%: IGF-binding protein complex acid-labile subunit (ALS, r = -0.654, p = 8.96E-05), IGF-binding protein 3 (IGFBP-3; r = -0.605, p = 3.98E-04) and IGF-2 (r = -0.601, p = 4.42E-04), shown in Figure 4.

**Figure 4.**
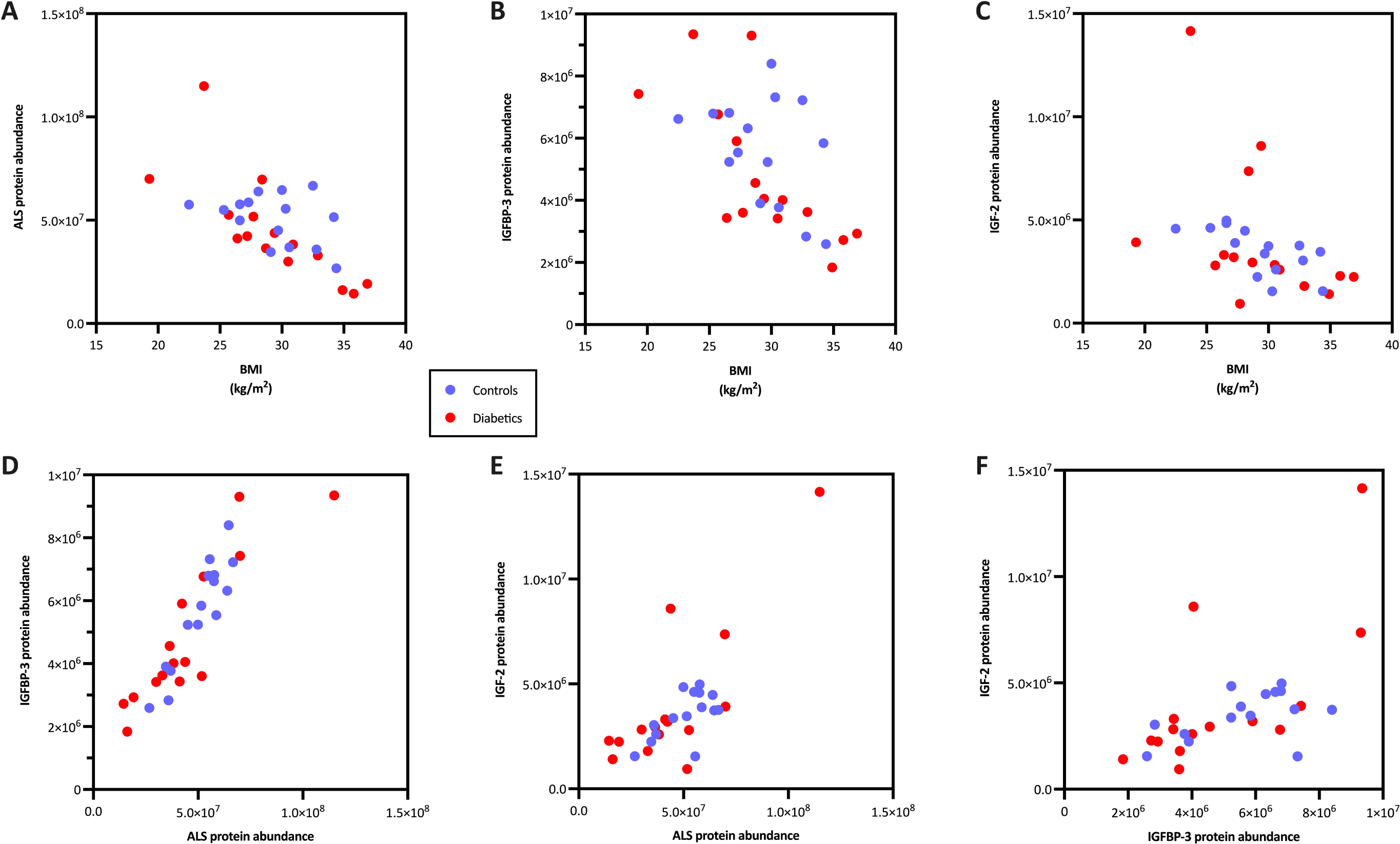
Significant correlations between BMI and components of the ternery complex composed of ALS (A), IGFBP-3 (B) and IGF-2 (C). Panels D through F show associations between ALS, IGFBP-3 and IGF-2. Data from control subjects shown in blue and patients with diabetes in red.

Remarkably, all three proteins have previously shown to be the components of a single plasma complex termed the ternary complex, as part of the insulin-like growth factor storage and transport mechanism in the circulation. Moreover, all of the correlations were negative, strongly suggesting that the levels of the ternary complex are reduced in the serum in obesity.

Analysis of individual peptides gave no further insight, and we found no specific PTMs that were associated with BMI after correction for multiple testing using FDR at 20%.

## Discussion

There is a strong need for new treatments for T2DM. This study aimed to identify differences between the serum proteome of individuals with T2DM and non-diabetic controls, to potentially provide insights into the aetiology of T2DM and identify novel targets for future research. While others have approached this question recently using proteomics, none have used a set of individual serum samples to account for person-to-person variability in measured biomarkers. This study used a protocol optimised for quantitative analysis of peptides and proteins, together with identification of PTMs, making no assumptions about which features of the proteome were important.

The key finding of this paper is that the proteome of patients with T2DM is extremely similar to that of their matched controls, so similar in fact that when considering only the overall abundance of each protein we find no differences at all between the two groups. This is across 175 proteins with abundances ranging across five orders of magnitude. Several proteins have average abundances that are widely different in the two groups, but in each case there is no consistency across the individual patient pairs, so the differences do not approach statistical significance. This highlights the benefit of running individual serum samples in biomarker studies – pooling samples as has been done previously in many proteomic studies of T2DM markedly reduces the power of the analysis (31,32).

When we extended the analysis to investigate fasting serum glucose, we identified a strong negative association between serotransferrin and glucose. This was unexpected as serotransferrin was one of the most abundant 7 serum proteins depleted by the MARS7 spin cartridges used to improve depth of coverage of the proteomics analysis. Therefore, it is not known if the difference we see is due to differences in serotransferrin concentrations in the original serum samples or in differences in depletion between subject groups. However, low serotransferrin has previously been described to be associated with both new onset diabetes (33) and end-stage renal disease in patients with T2DM (34), which suggests that the results in this study may reflect differences that pre-existed in the undepleted samples.

Three of the other four proteins whose correlations with serum fasting glucose remained significant after FDR correction have not previously been shown to be associated with glucose. However, there is a potential role of the innate immune system in diabetes (35) and consequently the association between higher levels of C1q subunit C and lower serum glucose is plausible. Moreover, in this analysis C1q subunit B also negatively correlates with glucose in this dataset (r = -0.436, p = 0.016) although C1q subunit A does not (r = -0.176, p = 0.351; unadjusted p values in both cases).

The strongest associations we identified in this study were between serum glucose concentration and hexose modifications of lysine residues on abundant serum proteins. These associations are strong, and all positive correlations, with higher modification fractions associated with both T2DM and higher glucose concentrations. The glycation of abundant serum proteins on lysine residues has been well known for decades (36), and such a result is therefore not unexpected. However, we believe that this is the first time that non-hypothesis-driven proteomics has identified this biomarker of hyperglycaemia, illustrating the reproducibility and power of the method used.

The modification mass change of +162 Da reflects the addition of a monosaccharide. It cannot be determined using MS which monosaccharide has reacted to form the glycation, but although glucose is not the most reactive monosaccharide it is the major carbohydrate in humans and it has previously been noted that most glycated proteins are glucose adducts (37). The +162 Da glycation is well known to be the product of the early stages of the Maillard reaction, forming from Amadori rearrangement of the Schiff base following nucleophilic attack of the monosaccharide. However, while the error tolerant search we used included the ability to detect a wide range of advanced glycation end products (AGE) that were present in the Unimod database used, we identified no products of the late Maillard reaction. Note that this analysis did not investigate the presence of cross-linked products of later reactions, so it is possible that the glycations we see were precursors to cross-links that were subsequently invisible to our analysis. Another possibility is that the serum proteins we have analysed are turned over sufficiently rapidly to only show the early stages of the Maillard reaction occurring.

In this study, we identified associations between glucose and glycations on the serum proteins α2-macroglobulin, apolipoprotein A1 and apolipoprotein A2. Many serum proteins have previously been found to be glycated in human serum (38). Many of those studies have investigated albumin, but we removed the majority of albumin from our samples prior to analysis. Alpha-2-macrogloblin has previously been shown to be glycated in serum (39) but specific glycated lysine residues have not previously identified. By contrast, residues that are glycated following treatment with glucose in vitro have been identified on apoA1 and apoA2 (40). However, none of those sites matched the lysine residues we found hexose-modified natively and correlating with fasting serum glucose. This is not due to lack of coverage in those regions of the proteins – peptides including all of the sites described by Kjerulf et al. were identified in this study. All of the proteins we found glycated have additional lysine residues that were not found modified. This may be due to the abundance of the modified peptides being below our sensitivity, or the structure of the protein impacting on the accessibility of individual lysines to glucose in solution. We also assume that glycation of other serum proteins would also be found, were the sensitivity of our method greater. Previous studies that have used enrichment to look for glycations in human serum have identified substantially more proteins glycated in vivo (41). We did not investigate the potential functional consequences of the glycation observed, although previous studies have done so (reviewed in 42).

In addition to the analysis of associations between T2DM or glucose and the serum proteome we also identified a strong negative association between BMI and IGF-2, IGFBP-3 and ALS. These three proteins are found in the ternary complex in plasma, which extends the half-life of IGFs in plasma from minutes to hours. IGF-1 was not detected in this study, but there is a considerable literature on the association between IGF-1 and BMI, most of which (but not all) shows a negative correlation, although that may differ by subject age (43). There is limited published data on IGF-2 and BMI, but previous studies are also consistent with our observation (44). Note that in contrast to the results from analysis of T2DM and glucose, where associations were with PTM fractions, these associations are with serum protein abundances.

In summary, we have used an optimised highly quantitative bottom-up LC-MS/MS proteomics protocol with a non-hypothesis-driven bioinformatics strategy to investigate differences in the proteome between patients with T2DM and matched controls. No protein-level differences were identified between the two patient groups, but more in depth analysis finds simple glycation of abundant serum proteins to be markedly different. The contrast with the results from the protein-level associations with BMI demonstrates the power of flexible, open proteomics protocols that investigate proteins, peptides and PTMs without prior assumptions about which features may be important in target identification or biomarker-related studies. In order to investigate peptides and PTMs it is important that the proteomics protocols used are optimised for precision, rather than detection of as many proteins as possible regardless of quantitative performance. Finally, it is also essential to analyse individual subject samples so that the results take into account person-to-person variability, which is obscured when pooled samples are used as is often the case in bottom-up proteomics analyses.

## Acknowledgements

The authors acknowledge the assistance of Mike Deery and the Cambridge Centre for Proteomics, for initial discussions and running the LC-MS/MS. and the assistance of Estere Seinkmane, RxCelerate Ltd., in reviewing the manuscript. We would also like to thank the excellent team at Papworth Hospital NHS Trust, in particular Dr. Peter Schofield and Dr. Sarah Clarke, for their role in recruiting patients into the MaGiCAD study.

## Author contributions

DEM and DJG designed the study. DEM, TS and AdG analysed data. All authors contributed to discussion. DEM wrote the manuscript. All authors reviewed and edited the manuscript. DEM is the guarantor of this work and, as such, had full access to all the data in the study and takes responsibility for the integrity of the data and the accuracy of the data analysis.

---

Supplementary Table 1. Association of peptide abundances with serum glucose. Only shown are the 39 peptides that that remained significant following FDR correction at 20%. Peptides are shown arranged by protein.

